# Persistent maternal memory, transient offspring effects: the temporal and sex-specific architecture of heritable heat-stress responses in zebrafish

**DOI:** 10.64898/2026.09.04.749453

**Authors:** Anastasiia Berezenko, Dragan Stajic, Simone Oberhaensli, Gary Delalay, Claudia Isabelle Keller Valsecchi, Heike Schmidt-Posthaus, Irene Adrian-Kalchhauser

**Author notes:** **Corresponding Author** Irene Adrian-Kalchhauser.

## Abstract

Environmental experiences can influence subsequent generations, yet little is known about the dynamics of such molecular memories across time, reproduction events, and offspring development. Here we provide a temporal map of the transmission of environmental information through egg provisioning and its phenotypic consequences in zebrafish. Specifically, we follow the effects of a 2-week heat wave across successive matings and two unexposed generations along a complete adult-to-egg-to-adult experimental trajectory, assessing gonad responses, maternal RNA, embryo survival, larval behaviour, adult stress resistance and liver transcriptomes.

We find that oocyte maternal RNA retained a pronounced heat-shock signature across several weeks and multiple spawning events, with selected functional components persisting into oocytes of the next generation. Somatic responses followed a fundamentally different trajectory: effects on liver transcription were sex- and timepoint-specific, with minor intergenerational signatures of energy metabolism and metabolic stress in males. Parental heat exposure also altered offspring survival, larval behaviour and thermal tolerance, with progressively weakened effects across successive reproductive events and across generations.

Our results reveal distinct temporal dynamics of environmental memory in the germline vs the developing and adult soma, with persistent molecular information in maternal provisioning coexisting with progressively fading offspring phenotypes. Such selective retention and loss of parental information align with theoretical predictions about the progressively declining value of inherited information, and may determine the dynamics of cross-generational population-level responses to extreme events.

## Introduction

The regulation of transcriptional networks enables organisms to rapidly fine-tune physiological and metabolic pathways in response to environmental change. Such environmental information has also been shown to persist across generations independently of DNA sequence. Examples include temperature-induced transcriptional states persisting for multiple generations in *C. elegans* (Klosin et al. 2017), paternal alarm-cue exposure altering sperm small RNAs and offspring behaviour in zebrafish (Ord et al. 2020), and combined heat and hypoxia exposure altering maternally deposited heat-shock proteins and increasing offspring thermal tolerance in zebrafish (Lim und Bernier 2023). These studies link to evolutionary models predicting that parental exposure to specific environmental stressors can modify molecular or phenotypic states in offspring and, under some conditions, prime them for similar future environments (Bonduriansky et al. 2012).

Non-genetic inheritance is ubiquitous and has been observed across the tree of life (Adrian-Kalchhauser et al. 2020; Jablonka und Raz 2009). The mechanistic basis of such inheritance varies from epigenetic mechanisms of transcriptional control (e.g., DNA methylation or chromatin conformation) to maternal RNA deposition during oogenesis (i.e., maternal effects; (Adrian-Kalchhauser et al. 2018; Perez und Lehner 2019). Its stability (i.e., persistence time) depends on the species (Frejacques et al. 2026; Klosin und Lehner 2016) as well as the extent and duration of the initial environmental input (Klosin et al. 2017) and the presence of contrasting information (Houri-Zeevi et al. 2021).

Regardless of the underlying molecular machinery, non-genetic inheritance can alter the rate and direction of evolutionary change (Day und Bonduriansky 2011; Bonduriansky et al. 2012). Its evolutionary consequences, however, depend on the dynamics, information value, and phenotypic effects of the inherited information (English et al. 2015). Understanding these dynamics is therefore critical. How long does environmental information persist within an organism, and what are the dynamics of its decay? How long are effects maintained across generations? Do memory dynamics differ between soma and germline? These questions are particularly relevant under climate change, where species persistence depends on the capacity of individuals and populations to respond rapidly to increasingly variable and extreme environments (Donelson et al. 2018; Harmon und Pfennig 2021). The acquisition, persistence and loss of environmental information may therefore help determine the capacity of populations to withstand rapid global change.

Fish are particularly sensitive to environmental perturbation. In zebrafish, acute heat exposure induces extensive transcriptional reprogramming affecting protein processing, signalling, metabolism, cell-cycle regulation and immune-related functions, alongside changes in energetic state such as glycogen content and lipid metabolism (Long et al. 2012). During development, prolonged warming can cause morphological abnormalities and pronounced changes in hepatic morphology and metabolism (Adzijovski et al. 2026), while thermal stress also affects the nervous system, altering brain protein expression, behaviour, oxidative balance, mitochondrial function and epigenetic state (Nonnis et al. 2021; Topal et al. 2025).

Importantly, these effects need not end with the directly exposed individual, and teleost reproductive biology provides a particularly plausible route for such effects to persist. Most species continuously generate new oocytes through self-renewing germline stem cells, and extensively provision them with maternal RNAs, proteins, lipids and nutrients from both ovarian and systemic sources (Lubzens et al. 2017; Liu et al. 2022; Beer und Draper 2013; Hiramatsu et al. 2015; Yan et al. 2017). Because this provisioning continues throughout oocyte growth, maternal environmental conditions can influence successive cohorts of eggs. Indeed, maternal warming alters egg transcriptomes in round goby, rainbow trout, stickleback and cod (Colson et al. 2019; Fellous et al. 2022; Adrian-Kalchhauser et al. 2018; Skjærven et al. 2024). Teleosts may therefore be particularly sensitive to the timing and persistence of thermal stress, because a single event can potentially influence multiple waves of developing germ cells and successive offspring cohorts. However, very little is known about the persistence of these molecular responses after the stress has ended, how they change across repeated reproductive events, and which components remain detectable in subsequent generations.

Here we explore the dynamics of information retention across timepoints and generations by exposing adult zebrafish to a transient heat challenge. We follow the effects through a unique combination of molecular and phenotypic readouts, from the exposed liver and gonads to offspring development from the individual oocyte across offspring survival, larval behaviour, adult physiology, and onward into the next generation. We resolve the persistence of environmental information in exposed parents and its consequences across offspring development. We show that an environmental perturbation leaves a remarkably durable and structured molecular memory in oocytes, persisting across multiple matings and for at least one month after exposure, while offspring phenotypes progressively fade and somatic transcriptional responses follow fundamentally different trajectories. Environmental information is therefore neither uniformly inherited nor simply diluted over time: it is selectively retained, transformed and lost across tissues, reproductive events and developmental stages.

## Material and Methods

### Fish husbandry and breeding conditions

Zebrafish (“AB/Tubingen” strain) were reared in a recirculation system (ZEBCARE B.V., Netherlands) in 1- and 5-L polycarbonate tanks (Tecniplast, Italy) to adulthood and maintained in a flow-through system in 9L glass tanks. Fish were maintained under standard laboratory conditions (28 ± 1 °C; 12/12-h light/dark cycle, 600–700 μs/cm conductivity, pH 7.5 ± 0.5, 10% of water exchange daily) at a density of ≤ 5 fish/L and under a twice-daily feeding regimen with dry food according to developmental stage (“Zebrafeed < 100µm” from 5 till 10 dpf; “100-200µm” from 10 till 15 dpf, “200-400 µm” from 15 till 35 dpf and “400-600 µm” from 35 dpf; Sparos I&D, Portugal) in the morning and Artemia spp. (ZEBCARE B.V., Netherlands) in the evening (Licitra et al. 2024). Fish were mated every 2 weeks after reaching maturity (from 3-4 months of age). Per breeding tank (Tecniplast, Italy), five males were randomly paired with five females from the same group in stagnant water overnight and returned to their source tank in the morning.

### Project setup and experimental timeline

140 treatment zebrafish (F0) were exposed to a raised water temperature of 34 °C for two weeks, while 140 control fish were maintained at the husbandry temperature of 28 °C. Each group was split into 2 replicates of 70 fish, with 35 females and 35 males in each replicate. The experiment was structured around timepoints T0 (before treatment), T1 (2 days after treatment), T2 (two weeks after treatment), and T3 (four weeks after treatment), and three generations F0 (exposed), F1 (not exposed), and F2 (not exposed; Figure 1). At T0, animals were mated to release any mature germ cells, and sampling was performed for gonad histology. At T1, T2, and T3, animals were mated, offspring were raised (F1) and then mated again (F2). Data collection thus extended vertically – along three timepoints of the exposed generation – and horizontally – across 1-2 generations (depending on data type). Intergenerational analyses focused on T1.

**Figure 1.**
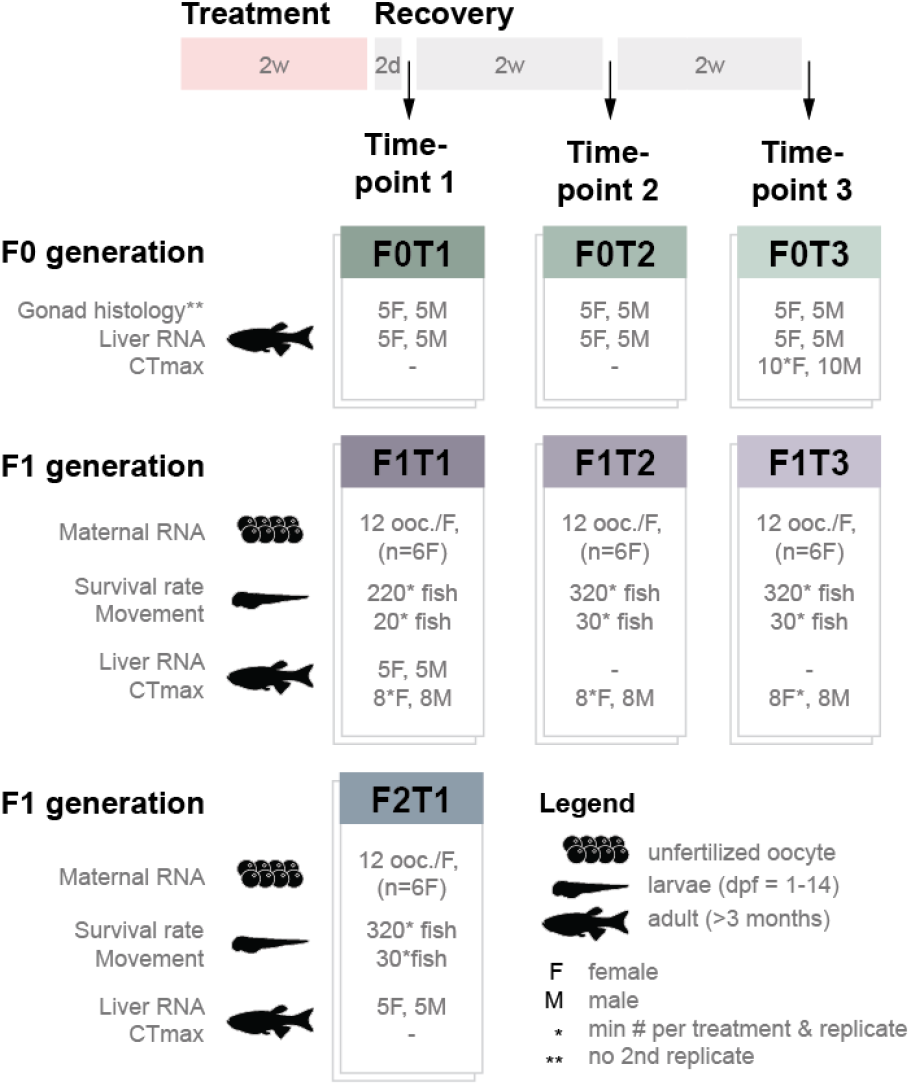
Experimental design and sampling scheme. Adult F0 zebrafish were exposed to a two-week heat treatment of 34°C or maintained under control conditions. This was followed by recovery at control temperature (28°C). Sampling and breeding were performed 2 days (T1), 2 weeks (T2), and 4 weeks (T3) after treatment. At each timepoint, F0 fish were analysed by gonad histology and liver RNA sequencing; Thermal tolerance (CTmax) was additionally measured at T3. In F1, we measured maternal RNA in unfertilised oocytes, larval survival and locomotor behaviour, liver RNA and CTmax as indicated. F1 fish derived from T1 were subsequently bred to generate the F2T1 generation, which was subjected to the same analyses. Numbers indicate the minimum numbers of females (F), males (M), oocytes (ooc.) or fish analysed per group replicate at each timepoint. For maternal RNA sequencing, 12 unfertilised oocytes were analysed per female from six females per group.

The following data types were collected:

1. gonad histology (F0T0, F0T1, F0T2, F0T3)
2. differential gene expression analysis of liver samples (F0T1, F0T2, F0T3, F1T1, F2T1)
3. differential gene expression analysis of unfertilized oocytes (maternal RNA) (F1T1, F1T2, F1T3, F2T1)
4. embryo survival rates (F1T1, F1T2, F1T3, F2T1)
5. larval movement upon stimulus (Danio Vision light-dark assay) (F1T1, F1T2, F1T3, F2T1)
6. adult temperature tolerance (CTmax assay) (F0T3, F1T1, F1T2, F1T3)

### Heat treatment

Both control and treatment groups were mated overnight to release accumulated mature germ cells and then transferred to four 9-liter glass tanks (2 replicates for control and 2 replicates for treatment, with 140 animals per tank). For two weeks, the control group was kept at 28 °C, while the treatment group was exposed to 34 °C using 300 W titanium heaters with temperature controllers (SCHEGO, Germany). The fish were monitored daily and fed as described before. No mortalities were recorded during this time. After two weeks, all tanks were set to 28°C for a two-day recovery period.

### Fish rearing

We raised F1 offspring of the exposed F0 parents at T1, T2, and T3, and the F2 offspring of non-exposed F1 sons/daughters at T1. To this end, fertilized eggs were collected from the mating tanks with a strainer and visually inspected. Developing embryos were transferred to Petri dishes (Cat No. 101, APTACA S.P.A., Italy) with E3 medium and methylene blue (0.5 mg/L) at a density of no more than 50 embryos per dish and incubated (BD series, Binder, Germany) at 28.0 °C under a 12:12 light cycle. To minimize the risk of disease, embryo surface sanitation was performed as described in the ZIRC protocol after 6 hpf (ZIRC). Offspring viability was checked daily, with dead embryos/larvae being recorded and removed from the dish, and the medium was exchanged daily. Food was provided from 5 days post-fertilization (dpf). At 14 dpf, juveniles were transferred from the Petri dishes into a recirculation system.

### Gonad morphology

To understand whether treatment affected offspring through alterations during germ cell development, we performed histology on the unexposed animals at T0 and exposed animals at T1, T2, and T3. To this end, 5 females and 5 males per timepoint and condition were euthanized with an overdose of buffered tricaine methanesulfonate ( > 0.3 mg/ml; Tricaine PHARMAQ 1000 mg/g, Pharmaq, Overhalla, Norway). and immersion-fixed in Davidson solution (Presnell et al. 1997) for 24 hours. Fish were sectioned longitudinally, dehydrated in a series of ethanol concentrations, embedded in paraffin, sectioned with a Microtome at 3 μm sections, and stained with haematoxylin and eosin (H&E). Slides were then digitalized with Nanozoomer S360 (Hamamatsu Photonics, Japan) with 0.23 µm/pixel resolution at 40x magnification and scored according to (Baumgartner et al. 2026a, 2026b). For females, cell annotation was performed manually in QuPath (v0.5.1) (Bankhead et al. 2017). For males, the slides were analysed in QuPath (v0.5.1), where first sperm cells were detected using the StarDist (v0.5.0) extension (Schmidt et al. 2018) and a pretrained model for brightfield H&E images (he_heavy_augment.pb; hematoxylin mean intensity used is above 0.5) and then annotated with a manually trained built-in object classifier (with at least 150 cells used for training each cell category). In females, previtellogenic (perinucleolar and cortical alveolar), vitellogenic (early and mid-/late vitellogenic), and mature oocytes were quantified manually according to the OECD Guidelines. In males, primary and secondary spermatocytes were classified as “spermatocytes”, while spermatids and spermatozoa were combined due to the inability of the segmentation model to reliably distinguish between these cell types. Spermatogonia were rarely detected by the segmentation model due to their lower haematoxylin intensity (< 0.5) and, when identified, were assigned to the “Other” category. The “Somatic cells” category comprised non-germline gonadal cells, including Sertoli and Leydig cells. The results were then analysed using R (v4.1.1) (R Core Team 2024) and RStudio Software (v2025.09.2) (Posit team 2025). For statistical comparisons of the cell proportions, a one-way ANOVA test with “BH” correction was used.

### Oocyte collection for RNA extraction

To understand whether treatment affected the information inherited by offspring, we performed DGE analysis of maternal RNA. To this end, we collected individual unfertilized oocytes of exposed F0 females at T1, T2, and T3, and of their unexposed F1 daughters at T1. Female fish were anesthetized with tricaine (in a concentration of 0.16 mg/ml; pH = 7) until gill movement slowed. Then, they were tapped dry with a paper towel, positioned laterally on their side in a Petri dish (Cat No. 101, APTACA S.P.A., Italy), and gently squeezed by applying light pressure with a finger on the lateral sides of the abdomen towards the urogenital opening. The Petri dish containing the oocytes was immediately placed on ice. The oocytes were then gently hand-picked under the stereomicroscope (SMZ745T, Nikon, Japan) with sterile forceps (Dumont #5, Dumont, Switzerland) and individually transferred to pre-cooled 1.5 ml Eppendorf tubes (Cat. No. 022431081, Eppendorf, Germany) with 100 ul DNA/RNA Shield (Cat. No. R1100-250, Zymo Research Corporation, USA). Once in preservative, oocytes were crushed with a sterile 20 µl pipette tip (Cat. No. 2149P-HR, Thermo Scientific™, USA) to ensure access of preservative to the oocyte contents. Tubes were then stored at −20°C until extraction.

### Offspring survival rates

To understand whether treatment in F0 affected the fitness of F1 and F2 offspring, we recorded the survival of larvae derived from exposed F0 parents at T1, T2, and T3, and from F1 parents at T1. The number of fish alive and dead was recorded between dpf 0 (after surface sanitation) and dpf 14 and assessed with Kaplan–Meier estimators (R package “survminer”; v0.5.0) (Kassambara et al. 2026).

### Movement upon stimulus (Danio-vision light-dark assay)

To understand whether treatment in F0 affected the behaviour of F1 and F2 offspring, we measured larval movement upon stimulus. To this end, we performed DanioVision light/dark treatments on F1 derived from exposed F0 parents at T1, T2, and T3, and of F2 offspring derived from unexposed F1 parents at T1. 5 dpf larvae were placed individually in the wells of the 96-well plate (Cat No. 92096, TPP, Switzerland) with 200 µl E3 medium (28 °C). The plate was then positioned in a DanioVision observation chamber (Noldus Inc., Netherlands) and exposed to a light-dark protocol as previously described (Ernst et al. 2023)). Specifically, a 40 min acclimatization in the dark was followed by 6 cycles of 10 min dark and 10 min bright phases. This protocol was first conducted at 28 °C, and then with another batch of naïve larvae at 34 °C. Larval movement was recorded at 1 min intervals with the EthoVision XT™ software (Noldus Inc., Netherlands). Incorrectly tracked fish (e.g.,, dead fish or fish for which software tracking failure occurred) were excluded from further analysis after visual inspection of the software-recorded videos. Per timepoint, group, and trial temperature, at least 30 naïve larvae were recorded, with 2-8 larvae among them excluded from further analysis (the number of larvae recorded per assay, along with the number excluded and used for further modelling, is provided in Supplementary File S1). The software-generated output files were analysed using R (v4.1.1) (R Core Team 2024) and RStudio Software (v2025.09.2) (Posit team 2025). Specifically, the response variable (the distance moved (mm)) was modelled with a Tweedie error distribution and a log link function using a generalized linear mixed-effects model (GLMM) fitted with the glmmTMB package (v1.1.14) (Brooks et al. 2017). Fixed effects included experimental “Timepoint” nested within “Group” (“Timepoint/Group”, where group indicates whether the fish were exposed to the heat shock), “Phase” (light or dark), “Temperature” (used during assay; 28 or 34 °C), and interactions among these factors. A random intercept and random slope for “CycleN_original” (the cycle order number, ranging from 1 to 12) were included for each “Well” (or Individual fish) to account for repeated measurements within wells. The model design was specified as: distance_moved_mm ~ Timepoint/Group * Phase*Temperature + cycle_time10:Cycle_N:Phase + cycle_time10 + (1 + CycleN_original | Well). The regressions were performed separately in order to model the movement of larvae from exposed parents at T1, T2 and T3, as well as the movement of F1 larvae from exposed parents at T1 and F2 larvae from unexposed parents at T1. Full model results, including estimates for all fixed effects and variance components for random effects, are provided in Supplementary Files S2 and S3 (for both regression models, respectively). Estimated marginal means (EMMs) for Group were calculated within each combination of Timepoint, Phase, and Temperature using the emmeans package (v2.0.1) (Lenth und Piaskowski 2026). Pairwise comparisons between groups were performed using reverse pairwise contrasts, and p-values were adjusted for multiple testing using the Holm-Bonferroni method. Estimated marginal means, confidence intervals, ratios, and pairwise comparisons are presented on the response scale.

### Thermotolerance (CT-max) assay

To understand whether heat treatment of F0 affected heat tolerance of exposed animals and their offspring, we measured the CT-max assay as described by Morgan et al. (Morgan et al. 2018) in exposed F0 parents at T3, and of non-exposed F1 adult offspring at T1, T2, and T3. To this end, a 9L glass tank was equipped with an Anova PRO heater (Anova Applied Electronics, Inc, China) installed on the right side of the tank and set to a starting temperature of 28°C. On the left side of the tank, a breeding net was installed, where a maximum of 10 fish were placed simultaneously and allowed to acclimate for 10 minutes. A digital thermometer Inkbird (INKBIRD Tech. Co., China), with an accuracy of ±0.1 °C, was installed near the breeding net to continuously measure the water temperature in this area of the tank. A GoPro Black 11 (GoPro Inc., USA) was used to record all trials. Then, the flow-through in the tank was shut down, and the temperature in the glass tank was increased at a rate of 0.5 °C per 2 minutes, with water mixing performed by the heater. When fish exhibited LOE (loss of equilibrium; loss of the ability to maintain an upright swimming position, defined as uncontrolled and disorganized swimming for two seconds), they were moved to an adjacent recovery tank at 28°C, and the water temperature during LOE, according to the thermometer, was recorded. Data was then analysed using R (v4.1.1) (R Core Team 2024) and RStudio Software (v2025.09.2) (Posit team 2025). Specifically, the Wilcox test for statistical inference with Bonferroni correction was used for LOE comparisons. The number of fish recorded per assay is provided in Supplementary File S4.

### Liver sampling

To understand whether soma and germline react and remember heat stress similarly, and to monitor systemic effects of heat treatment, we sampled livers of exposed F0 parents for RNA-seq at T1, T2, and T3, of non-exposed F1 offspring at T1, T2, and T3, and of non-exposed F2 grandchildren at T1. Fish were euthanized with an overdose of tricaine (in a concentration > 0.3 mg/ml). Then fish were positioned on their side in a sterile Petri dish (Cat No. 101, APTACA S.P.A., Italy) and the skin with the underlying muscle tissue was cut on and along the belly from the operculum to the anal fin as in (Gupta und Mullins 2010). The liver was grasped with Graefe forceps (Fine Science Tools GmbH, Germany) and collected in sterile pre-cooled 1.5 ml Eppendorf tubes (Cat. No. 022431081, Eppendorf, Germany) with 400 ml PBS (pH 7-7.4), positioned on ice and immediately extracted.

### Oocyte RNA extraction

Oocytes were retrieved from storage at −20 °C in batches of up to 24 samples and thawed on ice for five minutes. Total RNA was extracted with the Direct-zol™ RNA MicroPrep protocol (Cat. No. R2062, Zymo Research Corporation, USA), with the recommended adaptation for biological samples in DNA/RNA Shield (addition of 1 µl of Proteinase K (Cat. No. D3001-2-20, Zymo Research Corporation, USA) to the sample before the RNA purification step, mixing by inversion and incubation at room temperature for 30 minutes). The elution was performed with 11 µl of DNase/RNase-Free water and RNA was stored at −80 °C freezer (900 series, Thermo Scientific™, USA) before sequencing.

### Liver RNA extraction

Liver samples were dissociated into single cells prior to extraction, with the aim of obtaining material suitable for both ATAC-seq and RNA-seq. To this end, 10 µl of Collagenase A (in concentration 1 mg/mL; Roche, Switzerland) and 1 µl of DNAse I (in concentration 6 U/µl; Cat. No. E1011, Zymo Research Corporation, USA) were added to the liver samples, thoroughly mixed, and incubated at 37 °C for 30 minutes and 200 rpm in thermomixer (MKR23, Hettich Lab, Switzerland) with vortexing for 15 seconds every 10 minutes. Then, samples were spun down in 5mL sterile tubes with a cell-strainer cap (Cat No. 352235, Falcon®, USA) in a centrifuge (5810R, Eppendorf, Germany) at 500 rmp for 5 minutes. The single cell suspension was then mixed, and 150 µl used for the RNA extraction in accordance with the Direct-zol™ RNA MiniPrep protocol (Cat. No. R2052, Zymo Research Corporation, USA). The elution was performed with 50 µl of DNase/RNase-Free water, and samples were stored at −80 °C freezer (900 series, Thermo Scientific™, USA) before sequencing.

### RNA Sequencing

The quantity and quality of the purified total RNA was assessed using a Thermo Fisher Scientific Qubit 4.0 fluorometer with the Qubit RNA BR & HS Assay Kits (Cat No. Q32855 and Q10211, Invitrogen™, USA) and an Advanced Analytical Fragment Analyzer System using a Fragment Analyzer RNA Kit (Cat No. DNF-471, Agilent Technologies, USA), respectively. Oocyte samples with RNA integrity number (RIN) > 7 and RNA concentration above the detectable threshold (4 ng/µl) were used for sequencing. Liver samples without signs of rRNA degradation on the Fragment Analyzer graphs were chosen for sequencing, resulting in three liver samples from females and three from males per replicate of each timepoint. Input RNA samples (generally 100ng for oocyte RNA and 300ng for liver RNA) were first depleted of ribosomal RNA using a RiboCop rRNA Depletion Kit for Fish Kit following the Lexogen User Guide: 241UG837V0100 (Cat No. 241, Lexogen, Austria). Thereafter, cDNA libraries were generated using a CORALL Total RNA-Seq V2 library Prep.kit with UDIs 12nt set A1-A4 (Cat No. SKU 176, Lexogen, Austria) according to the protocol for long insert sizes and with 15 PCR cycles for liver RNA samples and 15-17 PCR cycles for oocyte RNA samples (Lexogen User Guide: 171UG394V0111). The resultant cDNA libraries were evaluated using a Thermo Fisher Scientific Qubit 4.0 fluorometer with the Qubit dsDNA HS Assay Kit (Cat No. Q32854, Invitrogen™, USA) and an Agilent Fragment Analyzer with a HS NGS Fragment Kit (Cat No. DNF-474, Agilent Technologies, USA), respectively. Equimolar-pooled libraries were sequenced paired-end using a set-up of 163:12:12:151 using two Illumina NovaSeq 6000 S4 Reagent Kits v1.5 (300 cycles; Illumina, 20028312) on an Illumina NovaSeq 6000 instrument. The quality of the sequencing runs was assessed using Illumina Sequencing Analysis Viewer (Illumina version 2.4.7), and all base call files were demultiplexed and converted into FASTQ files using Illumina bcl2fastq conversion software v2.20. The quality control assessments, production of libraries, and sequencing were carried out at the Next Generation Sequencing Platform, University of Bern.

### Read preprocessing

The preprocessing of the RNA-seq data was conducted using the nf-core rnaseq (Ewels et al. 2020) pipeline (release 3.14.0), with STAR (Dobin et al. 2013) used as the aligner and Salmon (Patro et al. 2017) used for quantification. For annotation and alignment, the primary assembly for zebrafish was used (GRCz11, release 112) (Howe et al. 2013). The obtained read numbers per sample with their mapping statistics are listed in Supplementary Files S5 for liver data and S6 for oocyte data. After preprocessing the samples were investigated for the presence of any sequencing quality issues (e.g., high amount of secondary alignment, high percentage of mt-rRNA, etc). If any of those were identified, the samples were not included in the further analysis. The numbers of samples sequenced, excluded (with the reasoning behind), and further analysed are provided in Supplementary Files S7 for liver data and S8 for oocyte data.

### Differential expression analysis

Differential expression analysis was performed with the DESeq2 package (v1.44.0; (Love et al. 2014) in R (v4.1.1) (R Core Team 2024) and RStudio Software (v2025.09.2) (Posit team 2025). Genes with less than 10 reads for the smallest group size (the smallest number of oocytes per mother (n=6) or the smallest number of liver samples per group (n=5)) were excluded from further analysis. Genes were considered significant when the abs. log2 fold change was ≥ 2 and adjusted P-value ≤ 0.05 for oocyte data and the abs. log2 fold change was ≥ 1.5 and adjusted P-value ≤ 0.05 for liver data. For sample clustering analysis, variance-stabilizing transformation (vst) was performed. DESeq2 results are available in Supplementary Files S9 and S10.

### Expression modules

To understand co-regulation of biological processes or functions in the dataset, interaction networks were generated separately for each timepoint in oocytes for genes with an abs. log2 fold change ≥ 2 and adjusted p-value < 0.05, and for each of timepoints in liver for genes with an abs. log2 fold change ≥ 1.5 and adjusted p-value < 0.05. Node and edge tables were created using STRINGdb (package v2.16.4; database v12.0, species “7955”, score threshold = 400) in R and imported into Cytoscape (version 3.10.4). The timepoint-specific networks were then combined for oocytes and livers, respectively, using a union merge based on ENSEMBL IDs. Node size was scaled according to connectivity in the merged network.

### Data availability

The sequencing data for this study is deposited in the European Nucleotide Archive (ENA) at EMBL-EBI under accession number PRJEB124081.

### Code availability

The R and shell scripts to replicate the analysis and visualizations are available at FIWI-UniBe/Zebrafish_Heat-Shock_Inheritance.

## Results

We exposed 140 zebrafish individuals in two population replicates to 34 °C for two weeks. Subsequently, populations were transferred back to control 28 °C temperatures, and sampling of gonads and somatic tissue was performed at two-week intervals of recovery after heat shock (Figure 1). Furthermore, to assess the extent of transcriptional memory across generations, exposed individuals were mated every two weeks, and corresponding F1 and F2 generations were reared and maintained at 28 °C. No deaths due to heat exposure were recorded during the treatment.

### Female gonads resist heat exposure, while males show a transient disruption

To determine general effects of heat treatment on gonad morphology in the exposed parents, we examined female and male gonad histology before heat shock and at the three recovery timepoints after heat shock. We found that 2 days after treatment, at T0T1, male gonads are temporarily depleted of mature spermatozoa (75% vs 25% of cells; Figure 2E, F). However, by timepoint T2, two weeks later, they had recovered to control levels. We did not observe any statistically significant differences in cell abundances in female gonads nor in oocyte morphology at any timepoint (Figure 2).

**Figure 2.**
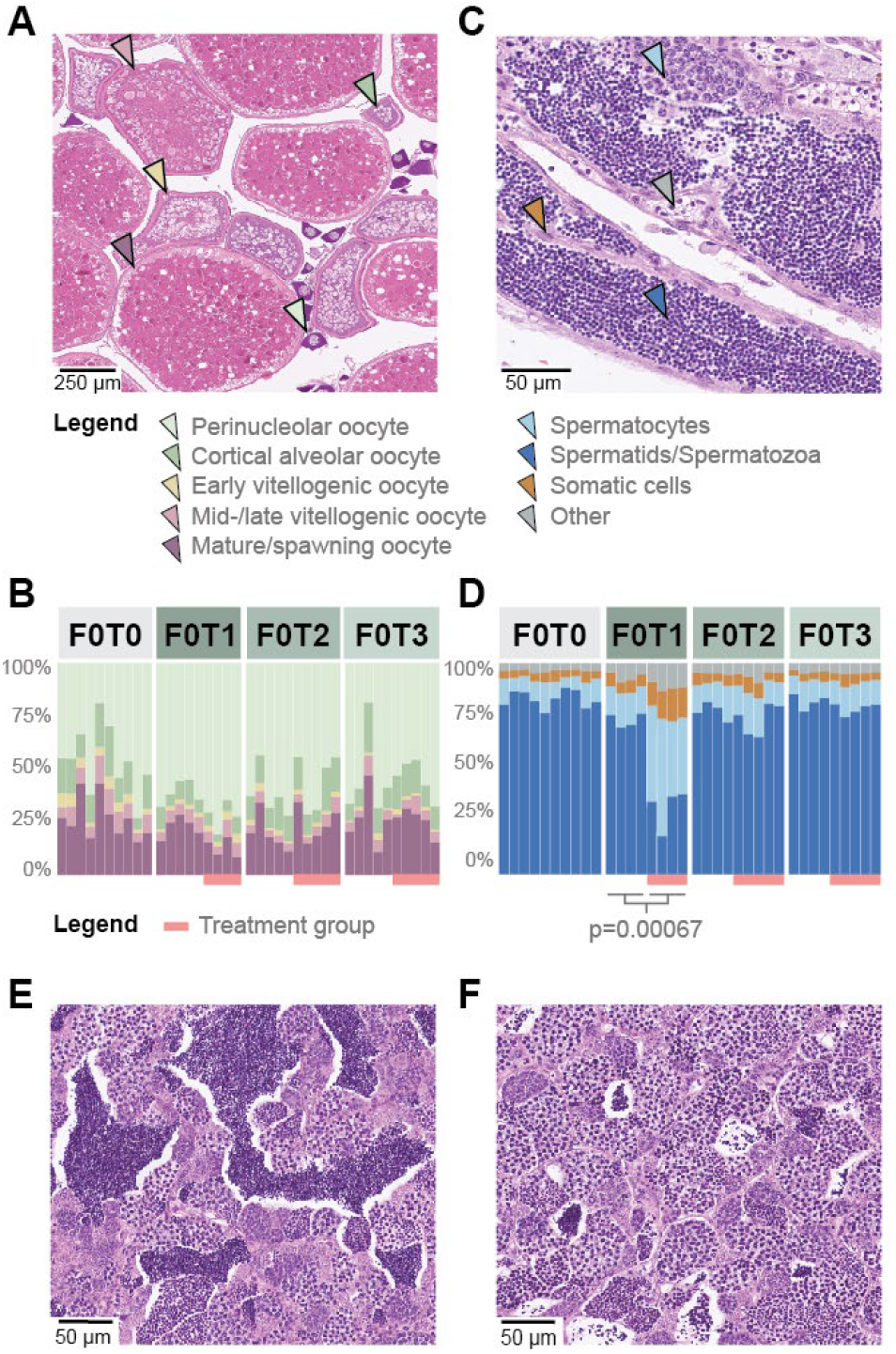
Heat exposure transiently reduces mature spermatozoa but does not alter ovarian morphology. **(A)** Representative ovary showing oocytes at different developmental stages (Perinucleolar, Cortical alveolar, Early vitellogenic, Mid-late vitellogenic, Mature). **(B)** Relative abundance of oocyte stages in female gonads before treatment (F0T0) and after treatment (F0T1–F0T3). Each bar represents one female; no treatment-associated differences were detected. **(C)** Representative testis showing successive stages of spermatogenesis (Spermatocytes, Spermatozoa) as well as somatic cells and other cells. **(D)** Relative abundance of male germ-cell stages in male gonads at F0T0–F0T3. Each bar represents one male. Mature spermatozoa were depleted in treated males at F0T1, two days after treatment, from 75% to 25% relative abundance, but were recovered by F0T2, two weeks after treatment. **(E, F)** Representative testes from control (E) and treated (F) males at F0T1, illustrating the transient depletion of mature spermatozoa two days after heat exposure.

### Oocytes retain a long-lasting heat-shock memory

To understand whether heat-shocked parents could pass information to their offspring through oocytes, we investigated the RNA content of mature, unfertilized oocytes as the first accessible timepoint of the next generations. Specifically, we sequenced the RNA content of mature, unfertilized, individual oocytes produced by exposed F0T1, F0T2, and F0T3 mothers and by unexposed F1T1 mothers. We determined differentially expressed genes (DEGs) by comparing mature oocytes of heat-shocked mothers and their daughters with corresponding oocytes from the control group. As oocyte production is continuous in zebrafish, with follicle growth and maturation occurring over approximately 8–10 days (Clelland und Peng 2009), oocytes collected at F1T2, F1T3 and F2T1 are unlikely to have been mature or late-stage oocytes during the heat treatment. In particular, oocytes sampled 30 days after exposure must have arisen from much earlier germ-cell stages, potentially including the germline stem-cell pool (Beer und Draper 2013).

We find that a total of 805 maternally provided RNAs react to heat shock when sampling oocytes from exposed mothers (F1T1: 382 up, 45 down, F1T2: 130 up, 57 down, F1T3: 117 up, 98 down at abs. log2 fold change >= 2 and adjusted P <=0.05; see Supplementary Table S10 for DESeq results). A candidate list of 56 genes known or predicted to be involved in the heat-shock response (Supplementary Table S11) was specifically examined, and indeed, five main components of heat-shock pathway were strongly upregulated in oocytes derived from exposed mothers (*serpinh1b, hsp90aa1*.*2, hsp70l, hspa8b* and *hsp70*.*2)*. Importantly, all five genes except *serbinh1b* remained elevated across all three successive F0 matings across 4 weeks after treatment (Figure 3E-I). Oocytes produced in the next generation by unexposed mothers lost this pattern, but still displayed differential gene expression between control and treatment (F2T1: 76 up, 87 down) (Figure 3D and 3E-I). This indicates a persistence of environmentally induced transcriptional changes in general, but a change in the exact composition in the response across generations (also compare (Adrian-Kalchhauser et al. 2020) Figure 3C).

**Figure 3.**
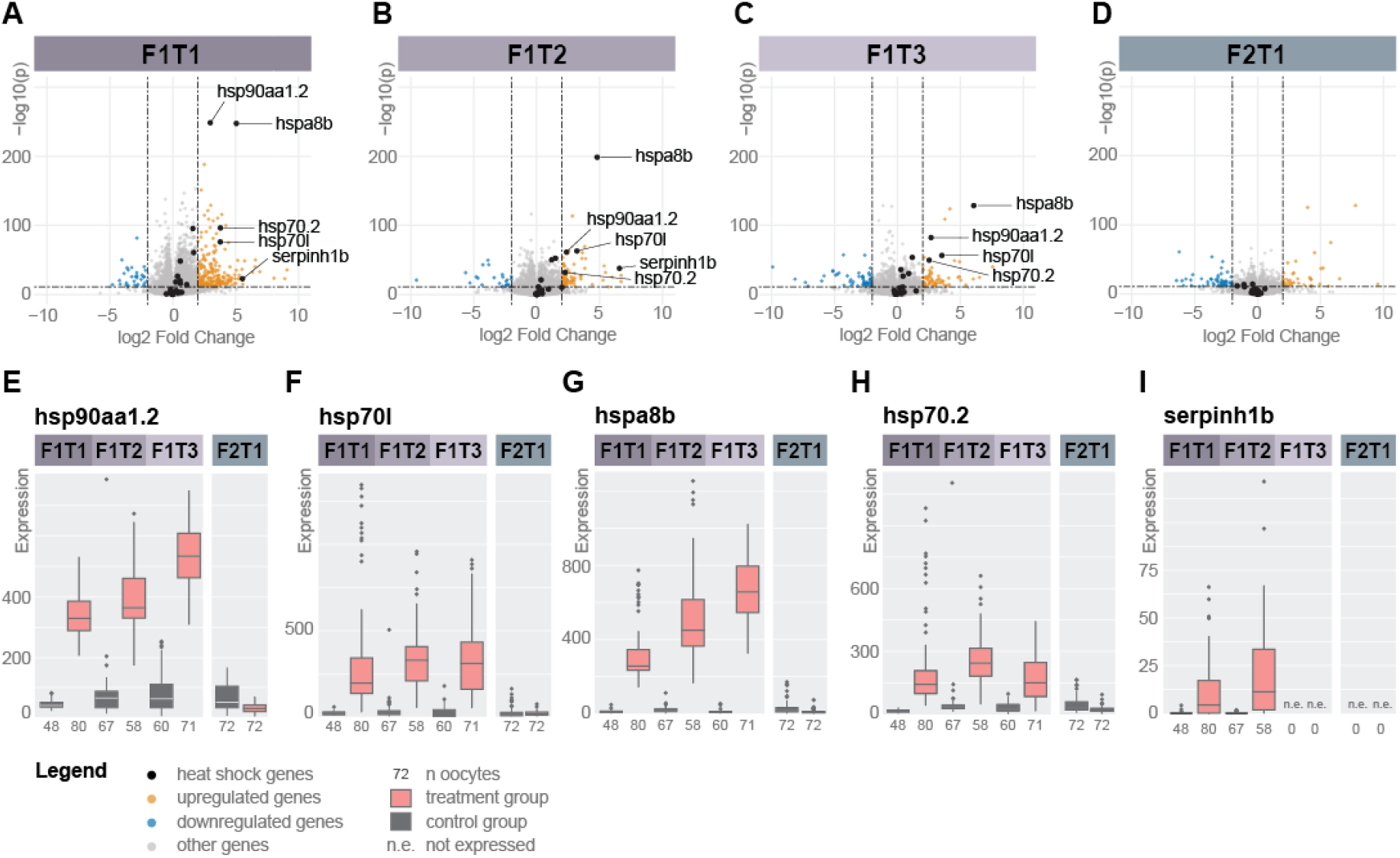
Oocytes retain a persistent heat-shock signature across successive spawnings. **(A–D)** Volcano plots of differential gene expression between treatment and control oocytes at F1T1, F1T2, F1T3 and F2T1. Upregulated genes are shown in orange, downregulated genes in blue and non-significant genes in grey (abs. log2 fold change ≥ 2, adjusted P ≤ 0.05). Heat-shock genes are highlighted in black and labelled when significantly regulated. **(E–I)** Expression of hsp90aa1.2 (E), hsp70l (F), hspa8b (G), hsp70.2 (H) and serpinh1b (I) across timepoints and generations in control (grey) and treatment (red) groups. For all genes except serpinh1b, expression remained elevated through F1T3, four weeks and three spawnings after treatment, but was no longer elevated at F2T1. Points represent individual oocytes; “n.e.” indicates not expressed.

### Oocytes retain structured memories of heat exposure across matings and generations

To further extend our understanding of maternal RNA regulation upon heat exposure, we performed network analyses on genes that were significantly up- and downregulated (adjusted P <= 0.05; abs. log2 fold change >= 2). 187 of all 805 differentially regulated genes formed an interaction network with modules related to heat response, circadian regulation, detoxification, immune signalling, lipid metabolism, and cytoskeletal organisation (Figure 4A). Within this network, several nodes were differentially regulated across timepoints or generations (Figure 4B-E). Within these networks, 18 genes were recurrently regulated in F1 oocytes, but not in F2, with 13 as part of the network and 5 outside of the network. Prominent network nodes that were consistently regulated across successive spawnings include the heatshock genes *hsp90aa1*.*2, hsp70*.*2, hspa8b, and hsp70l*, the apolipoprotein *apoeb* and the molecular chaperone *clu* from the lipoprotein module, the integrin beta *itgb8* from the cytoskeletal module, the transaminase *agxta* from the detoxification module, the metallopeptidase *mmp13a*.*2*, the sulfotransferase *sult2st3*, the nuclear receptor *nr1d2a*, the ubiquitin ligase regulator *trim109*, and the amino acid transporter *slc43a1b*. 60 nodes were regulated across generations, and 5 genes were regulated at all timepoints, 1 as part of the network and 4 as unlinked genes. These five genes include the embryonically expressed kinesin *kif26ab*, the ortholog of human FAM167B *fam167b*, the MHC class II protein complex member *mhc2d8*.*46a*, and two genes of unknown function (ENSDARG00000092702 and ENSDARG00000094960). The full list of significantly regulated genes is available as Supplementary Table S12.

**Figure 4.**
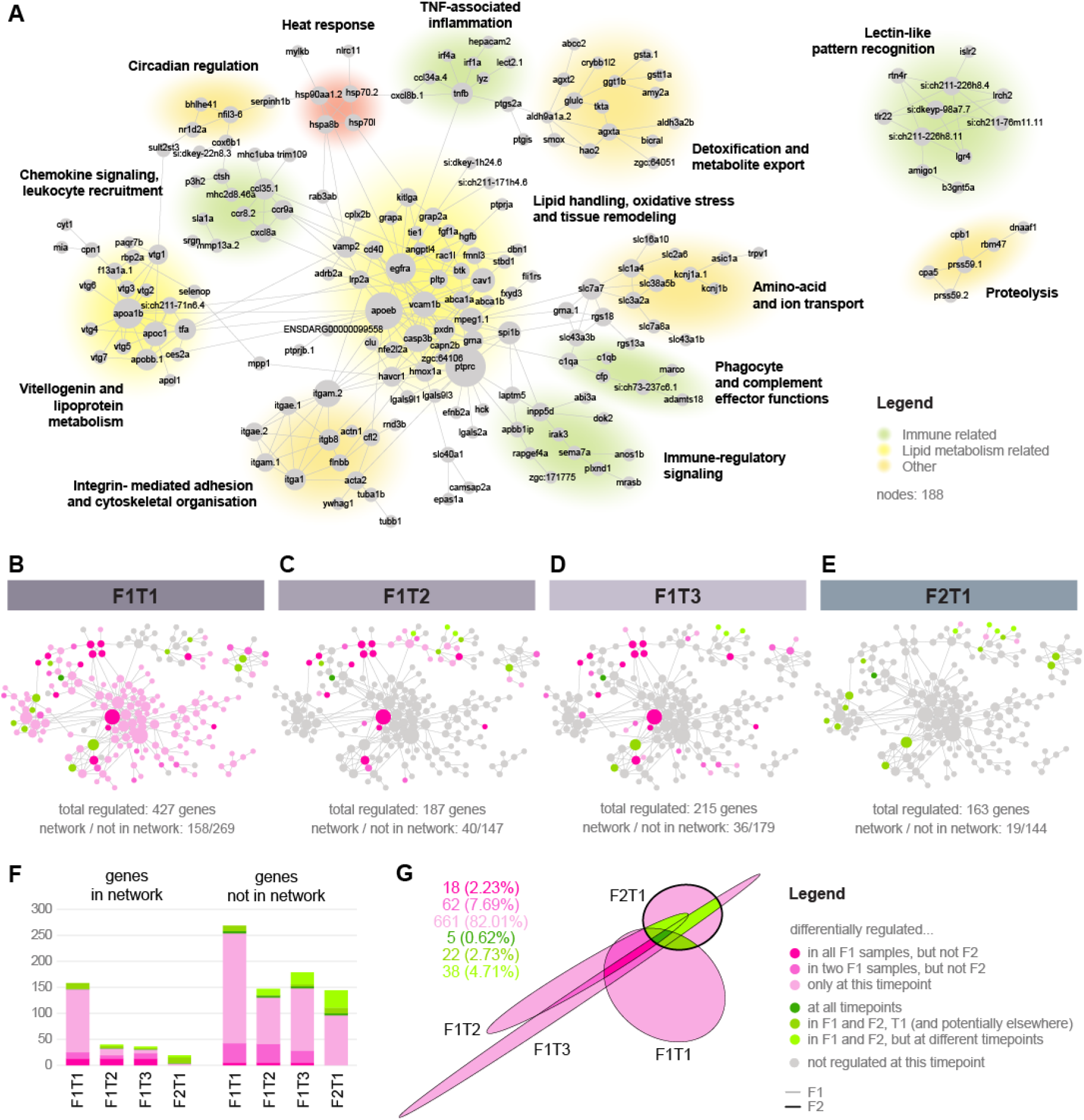
Oocyte responses progressively narrow while retaining recurrent functional components across time and generations. **(A)** Functional network of genes regulated in oocytes at one or more timepoints. Nodes represent genes, edges functional associations and node size connectivity; major functional modules are highlighted and labelled, and include a heat response module, five immune associated modules, three cytoskeleton and transport modules, two lipid metabolism modules and one proteolysis module. **(B–E)** Genes regulated at F1T1, F1T2, F1T3 and F2T1 mapped onto the network. Grey nodes are not regulated at the respective timepoint. Dark pink indicates recurrent regulation in F1, while green shades indicate recurrent regulation across generations. Grey nodes indicate genes not regulated at this timepoint. Numbers below each panel indicate total regulated genes and the numbers inside/outside the network. **(F)** Numbers of regulated genes inside and outside the network at each timepoint. **(G)** Graphical representation of overlaps between timepoints (proportional euler diagram). About 10% of genes are recurrently regulated in F1 (darker pink shades), and >7% display recurrent regulation across generations (green shades).

When examining the temporal patterns across the interaction network, we find a picture of a fading memory, with most network members regulated at T1, progressively fewer nodes regulated across T2 and T3 as well as in the next generation, and hardly any nodes regulated de novo beyond T1 (Figure 4B-E). The most prominently maintained module across spawning events was the heat shock module. Evidence for heritable regulation was most prominent in modules related to vitellogenin production, cytoskeleton, detoxification, and the immune modules lectin recognition, TNF associated inflammation, and chemokine signaling. Genes that were not part of the interaction network were less frequently regulated across all T1 timepoints, and more often regulated just at the investigated timepoint (Figure 4F).

### Phenotypic analyses reveal fitness-relevant effects in offspring embryos, larvae, and adults

Maternal RNAs play a profound role in directing early development in zebrafish. To assess whether the transcriptional changes had any observable phenotypic effects, we examined fitness-related traits such as likelihood of survival and thermal tolerance, as well as behavioral patterns.

We found that offspring embryo survival was compromised in F1, but not F2, with strongest effects at T1 (less than 50% chance of survival to adulthood, P value <0.0001, Kaplan-Meier estimator) and within the first 24 hours after fertilisation (Figure 5 A-D). While the probability of survival notably increased at subsequent time points (around 70%), it was still considerably reduced compared to control in T2 and T3 (P value <0.0001, Kaplan-Meier estimator).

**Figure 5.**
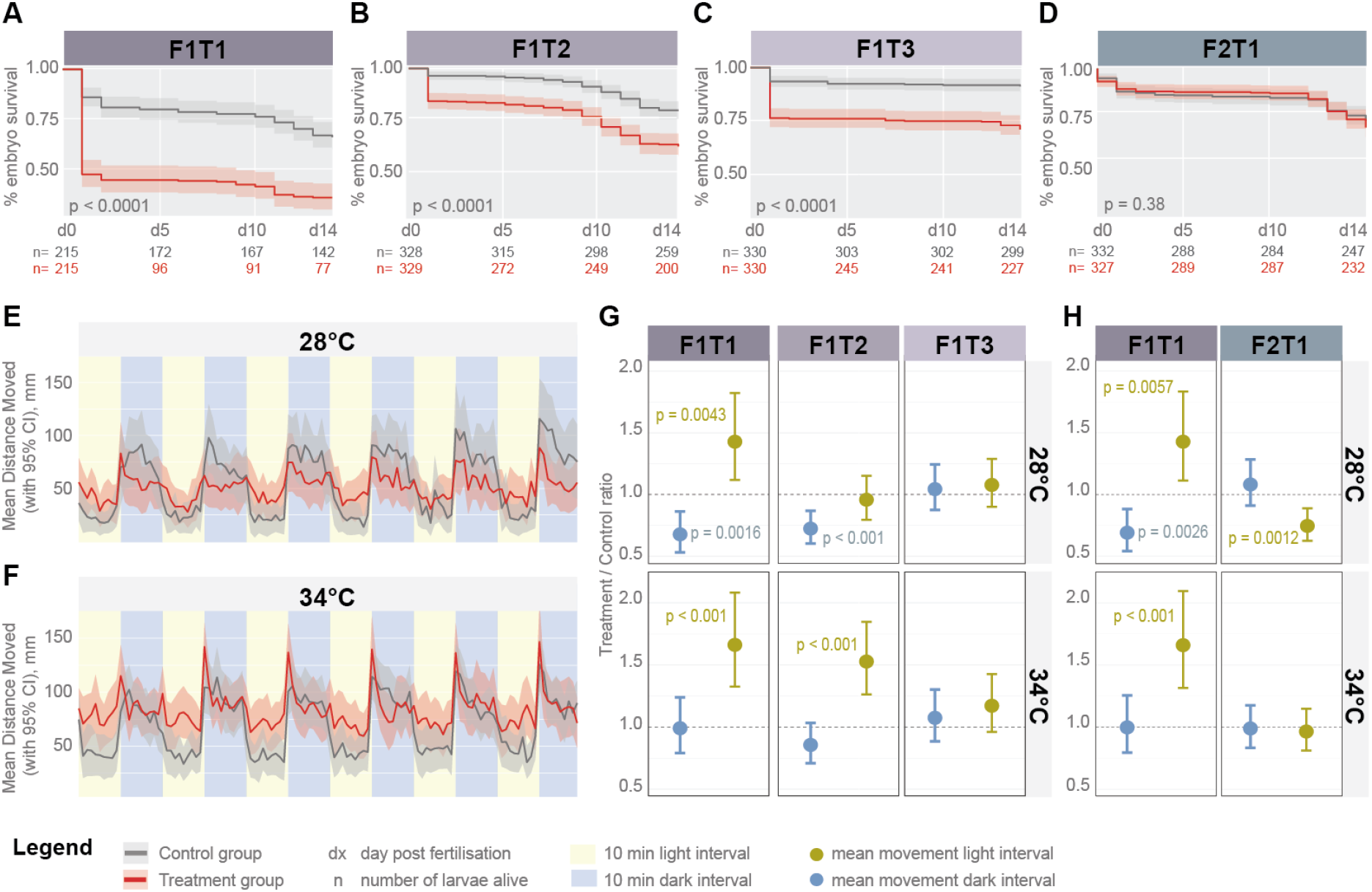
Parental heat exposure transiently reduces offspring survival and larval behaviour. **(A–D)** Larval survival from fertilisation to 14 days post-fertilisation (dpf) at F1T1, F1T2, F1T3 and F2T1. Grey and red lines show offspring of control and heat-treated parents, respectively; shaded areas indicate 95% confidence intervals. Numbers below each panel give the number of larvae alive at each timepoint. Survival was reduced in F1 offspring, with the strongest effect at T1, but not in F2. **(E, F)** Representative activity profiles of F1T1 larvae at 28 °C (E) and 34 °C (F) during alternating light and dark phases. Larvae from heat-treated parents (red lines) show a diminished light–dark response in comparison to control (grey lines). Yellow and blue backgrounds indicate 10-minute light and dark cycles, respectively. **(G, H)** Treatment-to-control ratios of larval movement during dark (grey) and light (yellow) phases across timepoints and temperatures. Values below 1 indicate reduced movement in offspring of heat-treated parents, and values above 1 indicate increased movement. P-values are included for timepoints with movement significantly different from controls. Separate models were used across same-generation F1 timepoints (G) and across two-generation T1 timepoints (H) The behavioural effect is evident at F1T1 and F1T2, but fades by F1T3 and is absent in F2T1.

As a behavioural phenotype, we monitored the response of 5 dpf larvae to the established light-dark assay (DanioVision) at 28 and 34°C. In this assay, larvae are exposed to repeated 10-minute alterations of light conditions, and tend to remain immobile during light phases and resume activity during dark phases (Burgess und Granato 2007; Hillman et al. 2024). While control animals followed the expected pattern of low movement in light and high movement in dark, treatment larvae displayed a more uniform response. They were moving significantly more in the light phase and were unable to maintain the same level of movement in the dark phase, despite an initial 1-minute movement spike at 34°C (Fig. 5E-H), and essentially failed to appropriately adapt their behaviour to the prevailing light condition. Similar to survival data, this behavioural effect faded across successive matings: at 28°, it was strongly detectable only at T1 and in the dark at T2; at 34°C, it was detectable at T1 and T2. It could no longer be detected at T3 or in F2.

The transfer of environmental information to subsequent generations can prime offspring for similar conditions and thereby improve performance when parental and offspring environments match (Bonduriansky et al. 2012; English et al. 2015). To assess whether parental exposure to heat increased thermal tolerance of their offspring, we used the CTmax assay. CTmax is determined by progressive heating until fish experience a loss of the righting reflex and thus the ability to swim upright. The exposed adults themselves did not show a significant increase in heat tolerance overall (41.53 °C ± 0.47 vs 41.44 °C ± 0.32; p-value = 0.233), but formed a bimodal distribution of tolerated temperatures (Figure 6). This might suggest a diversified, bet-hedging-like response, in which variation among individuals prepares the population for a range of possible future conditions. Such diversification could increase population persistence when environments are uncertain or fluctuate over time. In line with previous results from larvae, parental heat treatment increased thermal tolerance in offspring at T1 (41.50 °C ± 0.42 vs 41.25 °C ± 0.44; p-value = 0.032) and T2 (41.08 °C ± 0.42 vs 40.88 °C ± 0.48; p-value = 0.043), but not T3 (41.19 °C ± 0.52 vs 41.16 °C ± 0.54; p-value = 0.69; Fig. 6). Thus, parental heat exposure produced a transient increase in offspring thermal tolerance, while the directly exposed generation showed increased phenotypic heterogeneity rather than a uniform shift in tolerance.

**Figure 6.**
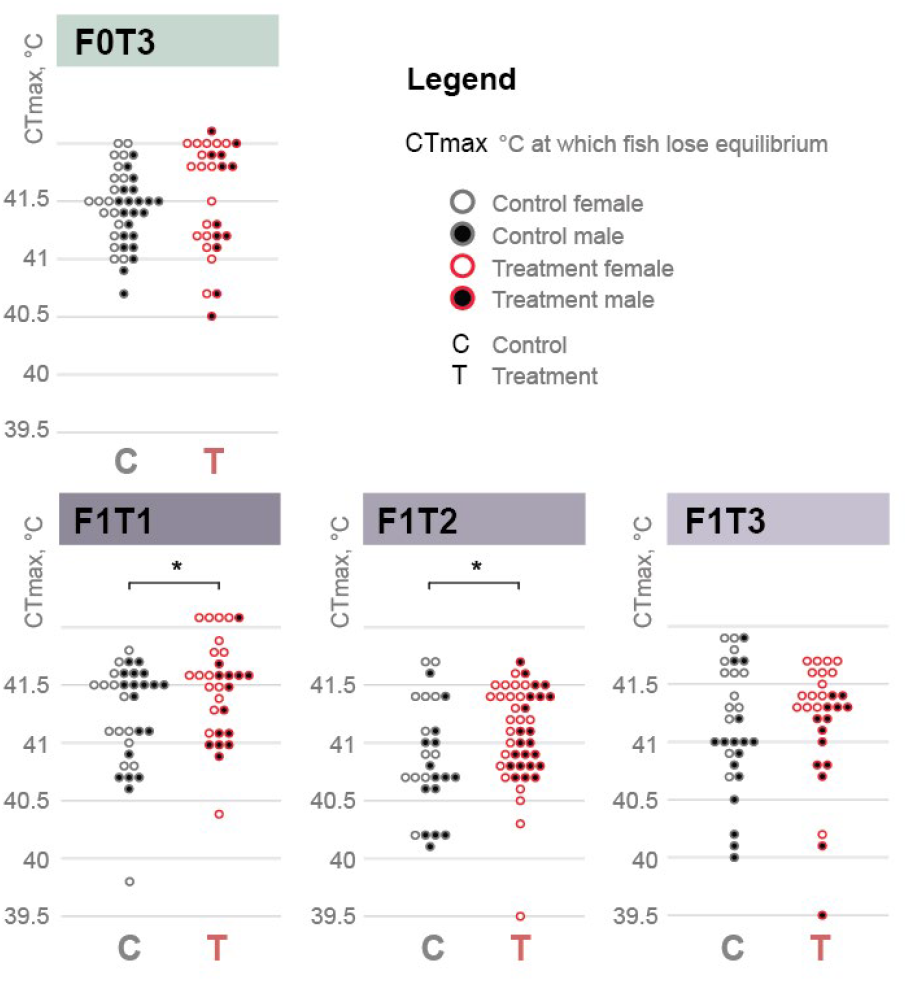
Parental heat exposure transiently alters offspring thermal tolerance. Critical thermal maximum (CTmax) was measured in directly exposed F0 adults at T3 and in unexposed F1 offspring at T1–T3. Each point represents one fish; grey and red indicate control and treatment individuals, while open and filled circles represent females and males, respectively. Mean CTmax was unchanged in F0, although treated fish showed a bimodal distribution. F1 offspring from heat-treated parents displayed a significantly higher CTmax at T1 and T2, but not at T3. Asterisks indicate significant differences after multiple-testing correction.

### Liver responses differ fundamentally from oocyte memory

Multicellular organisms are characterized by a clear developmental separation of germ and somatic cell lines. To understand whether these tissues with very different roles, and distinct tasks regarding inheritance, nonetheless mount similar responses to a systemic challenge, we characterized the gene expression in livers of exposed males and females, their offspring, and their grandchildren (F0T1-T3, F1T1, and F2T1). Livers were chosen as a central metabolic and stress-responsive organ that integrates systemic physiological signals. Overall, 614 genes were differentially regulated across the experimental timeline between treatment and control (the DESeq2 results are provided in Supplementary Table S9). Females and males mounted completely different responses, with 127 and 5 genes regulated in males or females respectively at F0T1, 122 and 5 at F0T2, 28 and 183 at F0T3, 97 and 33 at F1T1, and 90 and 44 at F2T1. Overlap between sexes was minimal, with no shared genes at F0T1 and F0T2, 4 shared genes at F0T3 (the apoptotic regulator *bbc3*, the predicted transcription factors *klf9* and *csrnp1a*, and the carboxylic acid transporter *slc16a9a*), and 1 shared gene at F1T1 and F2T1 respectively (the cytokine receptor cofactor *cisha* and the receptor tyrosine kinase *erbb3b*; Figure 7A). Most notably, 56 heat shock genes (Supplementary Table S11) were not among the significantly regulated genes (Figure 7B).

**Figure 7.**
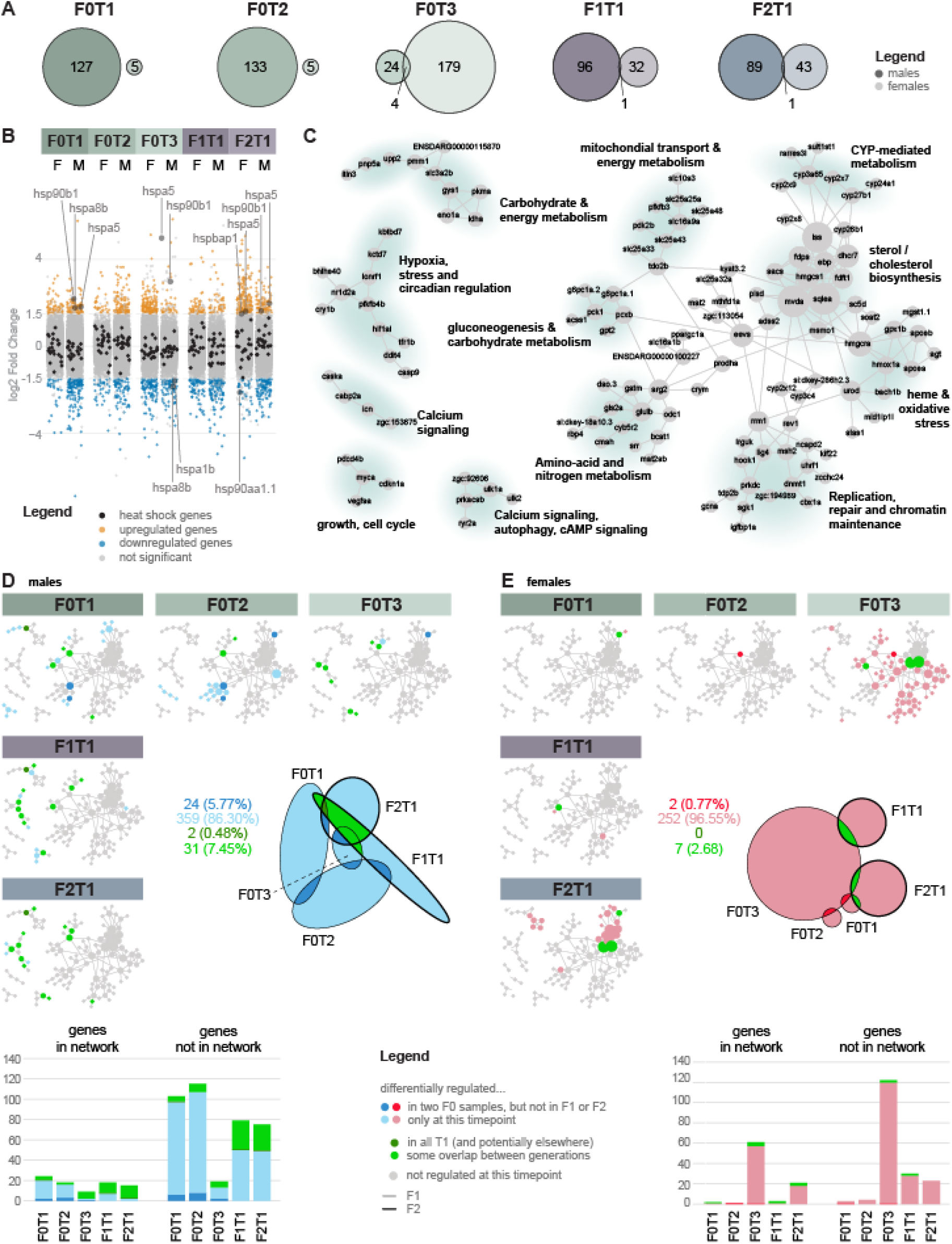
Liver transcriptome responses are strongly sex-, timepoint-, and generation-specific. **(A)** Numbers of differentially regulated genes per timepoint and sex. The overlap between males and females is zero in F0T1 and F0T2, 4 genes (2%) in F0T3, and 1 gene (0.79%) in F1T1 and 1 gene (0.76%) in F2T1. **(B)** Differential expression of heat-shock genes. Upregulated genes are shown in orange, downregulated genes in blue, and non-significant genes in grey; 56 heat-shock genes are highlighted in black. None are significantly or consistently regulated. **(C)** Functional network of genes regulated in liver. Nodes represent genes, edges represent functional associations, and node size represents connectivity; major metabolic, stress-response, and genome-maintenance modules are highlighted and labelled. **(D, E)** Regulated genes mapped onto the network for males **(D)** and females **(E)**, together with the numbers of regulated genes inside and outside the network at each timepoint (barchart), and a graphical representation of overlaps between timepoints (euler diagrams). The majority of genes are regulated at a single timepoint (bright blue/pink). Few genes overlap between consecutive F0 timepoints (dark blue/red). Inherited regulation (green; F1 and F2 are indicated by bold lines in the euler diagrams) is mostly restricted to males. Grey network nodes are not regulated at the respective timepoint.

We proceeded with assembling an interaction network to understand the regulation of functions and processes. We identified 12 modules mostly related to sugar and lipid metabolism, calcium signaling, cell cycle and chromatin maintenance, as well as stress. In contrast to oocytes, no immune modules were discovered (Figure 7C). Modules of the network were sex specific. In males, three modules representing energy-metabolism pathways were recurrently regulated across all three generations, and hypoxia/metabolic-stress and autophagy modules emerged in the unexposed F1 generation and were inherited to F2 (Figure 7D). Two genes were regulated in male livers at all timepoints: the ion antiporter *slc26a5* and the predicted phosphomannomutase *pmm1*. In females, network components were regulated in a delayed and timepoint-specific fashion at F0T3 and F2T1 (Figure 7E), and distinct modules were affected compared to males at each timepoint, with no gene consistently regulated across all timepoints. Surprisingly, hardly any genes were differentially expressed in female livers immediately after heat exposure. This might be attributable to strong resilience of gene expression in female livers, or to strong inter-individual variability that precludes significance. A list of genes and their regulation at specific timepoints is provided in Supplementary Table S13.

Given the provisioning role of livers for oocytes, and the potential importance of oocyte maternal RNA contents for larval liver development, we tested for systematic overlaps in gene expression regulation between oocytes and livers. To this end, we correlated log2fold-change values in connected samples, e.g., livers with oocytes produced by the same generation (e.g., F0T1 livers and F1T1 oocytes), and oocytes with livers from the same generation (e.g., F1T1 oocytes and F1T1 livers). This analysis did not reveal any correlation beyond individual spurious genes, indicating that somatic and germ cell responses to thermal stress are independent (Supplementary Figure S1). A gene overlap matrix of significantly differentially expressed genes in oocytes and livers across time points and generations is provided in Supplementary Table S14.

## Discussion

Understanding temporal dynamics of environmentally induced physiological and molecular responses within and across generations is of paramount importance because of their implications for evolutionary biology and conservation biology (Harmon und Pfennig 2021; Donelson et al. 2018; Bonduriansky et al. 2012). Understanding which information is transmitted, and for how long, is pertinent to understanding population-level adaptation capacities and limits of sexually reproducing organisms. Our study offers an unbiased, global, and thorough view of inheritance of transcriptional, fitness, behavioural, and physiological responses to an extreme environmental event across soma and germ cells in a vertebrate organism.

Results reveal that while phenotypic consequences of heat exposure fade along offspring development, in germ cells, the molecular memory persists (Figure 8A). The fading influence of parental heat exposure on offspring phenotypes may reflect the declining value of inherited information about past environments relative to conditions in the present environment. Parental cues are most useful when they predict the offspring’s environment; as offspring develop and acquire direct information themselves, inherited cues should become less influential (McNamara et al. 2016; Shea et al. 2011; English et al. 2015). Additionally, continued expression of a heat-associated phenotype might be costly and could become maladaptive over time.

**Figure 8.**
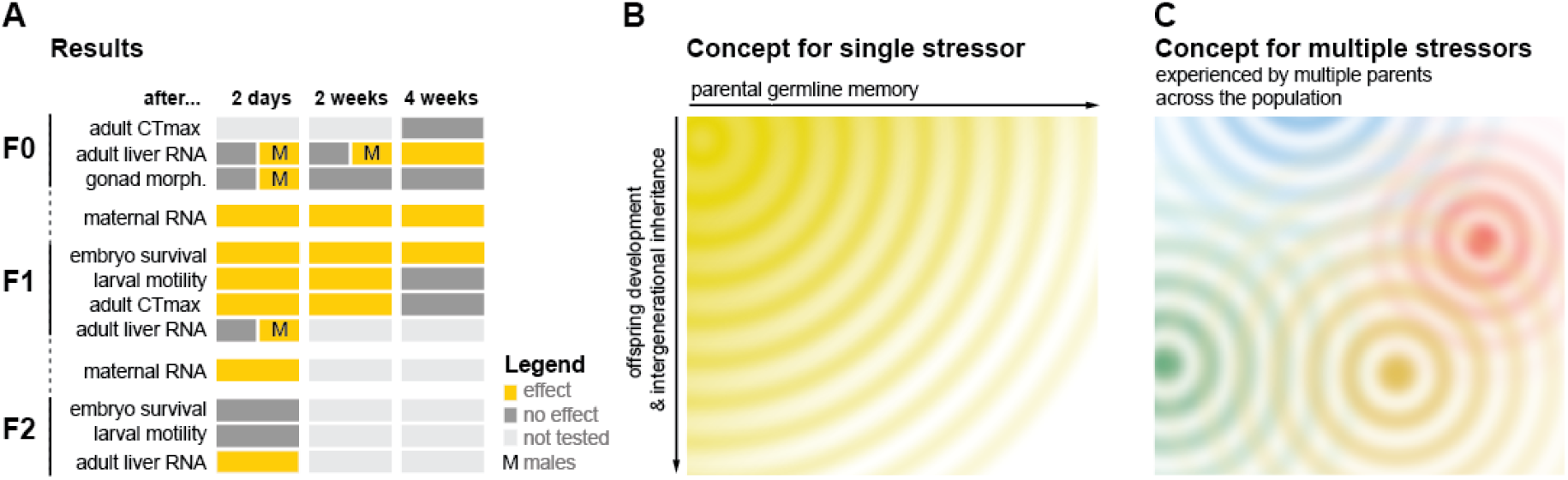
Summary and conceptual model of heat-stress memory across generations. **(A)** Summary of treatment effects across generations and readouts as observed in this study. Yellow indicates that an effect was observed, dark grey that no effect was observed, and light grey indicates timepoints or readouts that were not tested; “M” denotes male-specific effects. **(B)** Conceptual model for the fading of a single stress event: parental germline memory persists but with time is restricted to certain genes, molecular and phenotypic effects weaken across offspring development and generations. **(C)** Conceptual model for the interaction of stressors and their fading memories in natural populations: overlapping parental effects may generate dynamic and heterogeneous states and/or responses across time and individuals.

The strongest and most persistent molecular memory of heat exposure was found in the oocyte. While female gonads appeared morphologically unaffected by treatment, maternal RNA retained a clear heat-shock signature across repeated spawnings and during at least four weeks of recovery within the same generation, with little evidence of decay. Maternal RNA in F2 generation also was affected by the parental heat exposure, albeit not a classical heat shock response. Thus, a short, single environmental event can have significant influence not only within the same generation after the initial exposure, but also across generations. The broader transcriptional response narrowed over time, and only selected components of an identified interaction network persisted into the next generation. Early embryos of zebrafish, but also other non-mammalian vertebrates and invertebrates initially depend on maternally deposited molecules (Kojima et al. 2025), and perturbation of maternal factors can lead to substantial phenotypic effects during downstream development (Pikulkaew et al. 2011; Wagner et al. 2004). Our data suggest that maternal RNA may, in addition to its developmental roles, act as a long-term reservoir for a variety of environmental information, which is then weighed by offspring against true conditions in a decision framework integrating all available information about the environment (English et al. 2015; McNamara et al. 2016).

Where does the molecular memory reflected in the maternal RNA content of mature oocytes originate? Zebrafish follicle growth takes roughly ten days (Connolly et al. 2014; Clelland und Peng 2009). A heat signature present in mature oocytes 30 days after exposure therefore is not readily explained by exposed oocytes completing an altered maturation. The memory could, however, reside in early germ cells, including self-renewing germline stem cells (Beer und Draper 2013; Liu et al. 2022). An alternative candidate are tissues that provision the growing oocyte, such as the somatic germline or the liver. Follicle and theca cells continuously communicate with developing oocytes, and liver-derived vitellogenins and lipoproteins are systemically provided (Clelland und Peng 2009). However, our results show a relatively weak female liver response in contrast to the persistent memory in oocytes. Female zebrafish liver has previously been reported to be less transcriptionally responsive than male liver to perturbations (Zheng et al. 2013). Males, on the other hand, display some evidence for crossgenerational memory in selected pathways.

Taken together, our results show that somatic environmental memory is strongly sex-dependent. More importantly, different tissues clearly show different signatures of heat response and the extent of its memory. While our data show a clear pattern of maternal transfer of environmental information, our study falls short of understanding a potential sperm-mediated paternal contribution to the inheritance process (Ord et al. 2022; Ord et al. 2020; Irish et al. 2024)). Further studies are needed to investigate how stress and trauma alter sperm regulatory RNAs and offspring phenotype (Gapp et al. 2014; Rodgers et al. 2015; Chen et al. 2016).

The phenotypic changes observed in offspring behavior and thermal tolerance, together with persistent transcriptional patterns, are exciting candidates to mediate long-term implications on evolutionary dynamics. If an observed transcriptional memory modifies a phenotype on which selection acts, such as the ability of larvae to respond to a stimulus (which could be relevant to feeding efficiency or threat avoidance), even a short-term memory spanning one or two generations can conceivably affect the rate and direction of evolutionary change (Day und Bonduriansky 2011; Bonduriansky et al. 2012). Additionally, Heat exposure could potentially increase phenotypic diversity rather than shift all individuals in the same direction. Such diversification could provide the raw material for a bet-hedging strategy when future environments are unpredictable, although increased variance alone does not demonstrate bet hedging (Simons 2011; Proulx und Teotónio 2017). The single oocyte RNA sequencing data will allow us to explore the impact of parental stress on offspring variability on the molecular level in the future.

We conclude that environmental memory does not behave as a single inherited programme: its strength and persistence differ across readouts, molecular layers, sexes and life stages. A singular stress event thus generates a wave of altered offspring phenotypes across successive clutches and, potentially, generations, but the wave eventually dissipates if the environment remains at pre-event values (Fig 8B). In nature, a multitude of distinct stressors could generate overlapping “ripples” of information originating from different individuals and reproductive events, producing substantial temporal and spatial variation in available non-genetic information and, thus, phenotypes (Figure 8C).

## Acknowledgements

We are grateful to Xinyi Cheng, Kerim Klenja, Thea Baiardi, and Ramona Lüthi for taking great care of our zebrafish and the facility, to Denis Grandgirard from the Institute for Infectious Diseases for sharing and supporting the use of their Danio Vision equipment, to the Next Generation Sequencing Platform of the University of Bern for their support in planning and performing the high-throughput sequencing experiments for this paper, to Pei-Hsuan Wu from the University of Lausanne for advice and collaboration during attempts to isolate sperm RNA, and to Ursula Reinhart and Astrid Chanfon from the histology service group from the ITPA at the Vetsuisse Bern.

## Author contributions

AB - Design, Methodology, Data collection, Data Curation, Validation, Formal analysis, Writing (original draft preparation, review and editing), Visualization

DS - Conceptualization, Design, Methodology, Data collection, Data Curation (for Liver seq), Validation, Formal analysis, Writing (original draft preparation, review and editing), Visualization (Liver seq)

SO - Methodology, Writing (review and editing)

IAK - Conceptualization, Design, Data analysis strategy, Validation, Writing (original draft preparation, review and editing), Visualization, Supervision, Project Administration, Funding Acquisition

HS - Validation, Data Curation (histology), Writing (review and editing)

GD - Formal analysis

CVK - Experimental design, methodology

## Permits

Animal experiments described in this paper were carried out under permit BE18 of the Canton of Bern, Switzerland

## Funding

University resources and by funding of the Swiss National Science Foundation in the context of project grant #204838 supporting the integration of Ukrainian refugees into Swiss research groups.

## Data availability

See end of Methods section

## Notes

### Competing Interest Statement

The authors have declared no competing interest.

