## Supplementary material for "Persistent maternal memory, transient offspring effects: the temporal and sex-specific architecture of heritable heat-stress responses in zebrafish": SupplementaryFile01.TableS1_DanioVision_numbers.docx

**Supplementary File 01.**

**Table S1. Number of larvae used in the Movement upon stimulus (Danio-vision light-dark assay).**

Fish were tested at three different time points (T1, T2 and T3) across two generations (F1 and F2). Larvae obtained from exposed (treatment group) and unexposed (control group) parents were tested with the Danio Vision assay at 5 days post fertilization. Specifically, a 40 min acclimatization in the dark was followed by 6 cycles of 10 min dark and 10 min bright phases. This protocol was first conducted at 28 °C, and then with another batch of naïve larvae at 34 °C. Larval movement was recorded at 1 min intervals with the EthoVision XT™ software (Noldus Inc., Netherlands). Incorrectly tracked fish (for example, dead fish or fish for which software tracking failure occurred) were excluded from further analysis after visual inspection of the software-recorded videos.

| Time point (Population) | Group | Trial temperature | | | | | |
| --- | --- | --- | --- | --- | --- | --- | --- |
|  |  | 28 °C | | | 34 °C | | |
|  |  | N of fish  used in the trial | N of fish  analyzed | % of fish excluded | N of fish  used in the trial | N of fish analyzed | % of fish excluded |
| T1 (F1T1) | Control | 36 | 23 | 36,1 | 35 | 27 | 22,9 |
|  | Treatment | 31 | 23 | 25,8 | 31 | 25 | 19,4 |
| T2 (F1T2) | Control | 48 | 44 | 8,3 | 48 | 44 | 8,3 |
|  | Treatment | 44 | 39 | 11,3 | 39 | 31 | 20,5 |
| T3 (F1T3) | Control | 48 | 46 | 4,2 | 44 | 40 | 9,1 |
|  | Treatment | 40 | 39 | 2,5 | 34 | 31 | 8,8 |
| T1 (F2T1) | Control | 48 | 48 | 0 | 47 | 46 | 2,1 |
|  | Treatment | 48 | 47 | 2,1 | 48 | 48 | 0 |
