## Supplementary material for "Persistent maternal memory, transient offspring effects: the temporal and sex-specific architecture of heritable heat-stress responses in zebrafish": SupplementaryFile02.TableS2_DanioVision_modelling_F1.docx

**Supplementary File 02.**

**Table S2. glmTMB model output for Danio Vision analysis from the data with larvae at T1, T2 and T3 timepoints at F1 generation**

Fish were tested at three different time points (T1, T2 and T3) across two generations (F1 and F2). Larvae obtained from exposed (treatment group) and unexposed (control group) parents were tested with the Danio Vision assay at 5 days post fertilization. Specifically, a 40 min acclimatization in the dark was followed by 6 cycles of 10 min dark and 10 min bright phases. This protocol was first conducted at 28 °C, and then with another batch of naïve larvae at 34 °C. Larval movement was recorded at 1 min intervals with the EthoVision XT™ software (Noldus Inc., Netherlands). The software-generated output files were analysed using R (v4.1.1) and RStudio Software (v2025.09.2). Specifically, the response variable (the distance moved (mm)) was modelled with a Tweedie error distribution and a log link function using a generalized linear mixed-effects model (GLMM) fitted with the glmmTMB package (v1.1.14). Fixed effects included “Population” nested within “Group” (“Population/Group”), “Phase” (light or dark), “Temperature” (used during assay; 28 or 34 °C), and interactions among these factors. A random intercept and random slope for “Cycle” (the cycle order number) were included for each “Well” (or Individual fish) to account for repeated measurements within wells. The model output presented here (for fixed and random effects, respectively) is from modelling the movement of larvae from exposed parents at F1 generation at different timepoints T1, T2 and T3.

Fixed effect table

| **Term** | | | | **Estimate** | **SE** | **Statistic** | **Pvalue** | **Estimate_exp_** | **CI_exp_** |
| --- | --- | --- | --- | --- | --- | --- | --- | --- | --- |
| **Timepoint** | **Group** | **Temperature** | **Phase** |  |  |  |  |  |  |
| F1T1 | Control | 28 | Light | 3.185 | 0.090 | 35.541 | 0.000 | 24.172 | 20.278 - 28.814 |
| (Intercept) | | | |  |  |  |  |  |  |
| F1T2 | Control | 28 | Light | 0.058 | 0.109 | 0.534 | 0.593 | 1.060 | 0.856 - 1.311 |
| F1T3 | Control | 28 | Light | -0.024 | 0.109 | -0.216 | 0.829 | 0.977 | 0.789 - 1.209 |
| F1T1 | Control | 28 | Dark | 1.241 | 0.035 | 35.500 | 0.000 | 3.460 | 3.231 - 3.705 |
| F1T2 | Control | 28 | Dark | 0.336 | 0.041 | 8.258 | 0.000 | 1.399 | 1.292 - 1.515 |
| F1T3 | Control | 28 | Dark | 0.168 | 0.041 | 4.129 | 0.000 | 1.183 | 1.092 - 1.281 |
| F1T1 | Treatment | 28 | Light | 0.354 | 0.124 | 2.859 | 0.004 | 1.424 | 1.118 - 1.815 |
| F1T2 | Treatment | 28 | Light | -0.042 | 0.093 | -0.452 | 0.651 | 0.959 | 0.798 - 1.151 |
| F1T3 | Treatment | 28 | Light | 0.075 | 0.091 | 0.823 | 0.410 | 1.077 | 0.902 - 1.287 |
| F1T1 | Treatment | 28 | Dark | -0.738 | 0.046 | -15.893 | 0.000 | 0.478 | 0.437 - 0.524 |
| F1T2 | Treatment | 28 | Dark | -0.277 | 0.035 | -7.944 | 0.000 | 0.758 | 0.708 - 0.812 |
| F1T3 | Treatment | 28 | Dark | -0.032 | 0.034 | -0.946 | 0.344 | 0.969 | 0.907 - 1.035 |
| F1T1 | Control | 34 | Light | 0.466 | 0.119 | 3.917 | 0.000 | 1.594 | 1.262 - 2.012 |
| F1T2 | Control | 34 | Light | -0.183 | 0.148 | -1.235 | 0.217 | 0.833 | 0.623 - 1.113 |
| F1T3 | Control | 34 | Light | -0.725 | 0.150 | -4.840 | 0.000 | 0.484 | 0.361 - 0.650 |
| F1T1 | Control | 34 | Dark | -0.313 | 0.044 | -7.135 | 0.000 | 0.731 | 0.671 - 0.797 |
| F1T2 | Control | 34 | Dark | 0.090 | 0.054 | 1.666 | 0.096 | 1.094 | 0.984 - 1.216 |
| F1T3 | Control | 34 | Dark | 0.371 | 0.056 | 6.637 | 0.000 | 1.449 | 1.298 - 1.616 |
| F1T1 | Treatment | 34 | Light | 0.150 | 0.168 | 0.893 | 0.372 | 1.162 | 0.836 - 1.615 |
| F1T2 | Treatment | 34 | Light | 0.462 | 0.134 | 3.442 | 0.001 | 1.588 | 1.220 - 2.066 |
| F1T3 | Treatment | 34 | Light | 0.082 | 0.135 | 0.607 | 0.544 | 1.085 | 0.833 - 1.414 |
| F1T1 | Treatment | 34 | Dark | 0.224 | 0.060 | 3.713 | 0.000 | 1.252 | 1.112 - 1.409 |
| F1T2 | Treatment | 34 | Dark | -0.297 | 0.048 | -6.234 | 0.000 | 0.743 | 0.677 - 0.816 |
| F1T3 | Treatment | 34 | Dark | -0.054 | 0.051 | -1.058 | 0.290 | 0.947 | 0.857 - 1.047 |
| **Other** | | | | | | | | | |
| PhaseLight:cycle_time10:Cycle_N | | | | 0.003 | 0.001 | 4.058 | 0.000 | 1.003 | 1.001 - 1.004 |
| PhaseDark:cycle_time10:Cycle_N | | | | -0.005 | 0.001 | -8.119 | 0.000 | 0.995 | 0.994 - 0.996 |
| cycle_time10 | | | | -0.007 | 0.002 | -2.789 | 0.005 | 0.993 | 0.989 - 0.998 |

**Note**: ***Term*** – the name of the model fixed effect; ***Estimate*** – the estimated model coefficient on the log scale; ***SE*** – standard error of the estimate; ***Statistic*** – wald test statistic (z-value = Estimate/SE); ***Pvalue*** – wald test p-value; ***Estimate_exp_*** – the exponentiated coefficient; ***CI_exp_*** – lower and upper 95% confidence intervals for the exponentiated coefficient.

Random effect table:

| **Group** | **Term** | **Estimate** | **CI_exp_** |
| --- | --- | --- | --- |
| Well | sd__(Intercept) | 0.493 | 0.456 – 0.533 |
| Well | sd__CycleN | 0.046 | 0.042 – 0.050 |
| Well | cor__(Intercept).CycleN | -0.595 | 4.459 – 4.588 |

**Note**: ***Group*** – the grouping factor applied (here, “Well”, which is individual fish analyzed); ***Term*** – the name of the model random effect (the “Cor” = correlation); ***Estimate*** – the estimated model coefficient (here, variation among individuals); ***CI*** – lower and upper 95% confidence intervals for the term.
