## Supplementary material for "Persistent maternal memory, transient offspring effects: the temporal and sex-specific architecture of heritable heat-stress responses in zebrafish": SupplementaryFile04.TableS4_CTmax_numbers.docx

**Supplementary File 04.**

**Table S4. Number of adult fish used in the thermotolerance (CT-max) assay.**

Fish were tested with the CT-max assay as described by Morgan et al.^1^ at 3 different time points (T1, T2 and T3) and across two generations (F0 and F1). Specifically, exposed F0 parents at T3 and non-exposed F1 adult offspring at T1, T2, and T3 were analyzed.

| Time point (Population) | Group | Sex | |
| --- | --- | --- | --- |
|  |  | Female (n of fish tested) | Male (n of fish tested) |
| T3 (F0T3) | Control | 18 | 21 |
|  | Treatment | 16 | 12 |
| T1 (F1T1) | Control | 14 | 19 |
|  | Treatment | 16 | 14 |
| T2 (F1T2) | Control | 12 | 15 |
|  | Treatment | 22 | 22 |
| T3 (F1T3) | Control | 13 | 16 |
|  | Treatment | 15 | 14 |

^1^ Morgan R, Finnøen MH, Jutfelt F. CTmax is repeatable and doesn't reduce growth in zebrafish. Sci Rep. 2018;8(1):7099. doi:10.1038/s41598-018-25593-4
