## Supplementary material for "Persistent maternal memory, transient offspring effects: the temporal and sex-specific architecture of heritable heat-stress responses in zebrafish": SupplementaryFile07.TableS7_Outliers_Livers.docx

**Supplementary File 07.**

**Table S7. An overview of the sequenced liver samples.**

Before the differential gene expression analysis, the RNA-seq data obtained were investigated for the presence of any sequencing quality issues. If any of those were identified, the samples were not included in the further analysis. The table provides a detailed overview of the samples that are included and excluded (as well as the reasoning behind this).

| Timepoint (Population) | Group | Sex | Samples N | | | Excluded samples | Reason for exclusion |
| --- | --- | --- | --- | --- | --- | --- | --- |
|  |  |  | extracted | sequenced | analyzed |  |  |
| T1 (F0T1) | Control | Female | 10 | 6 | 6 |  |  |
|  | Control | Male | 10 | 6 | 5 | ML136 | Bad mapping results (the highest number of secondary alignments across all liver samples, overall bad mapping results) |
|  | Treatment | Female | 10 | 6 | 6 |  |  |
|  | Treatment | Male | 10 | 6 | 5 | ML146 | The mt-rRNA content of the sample exceeded 10% of the total amount of reads |
| T2 (F0T2) | Control | Female | 10 | 6 | 5 | FL211 | Separate clustering on the sample-to-sample distance matrix from all other samples, including both females and males |
|  | Control | Male | 10 | 6 | 6 |  |  |
|  | Treatment | Female | 10 | 6 | 6 |  |  |
|  | Treatment | Male | 10 | 6 | 6 |  |  |
| T3 (F0T3) | Control | Female | 10 | 6 | 6 |  |  |
|  | Control | Male | 10 | 6 | 6 |  |  |
|  | Treatment | Female | 10 | 6 | 6 |  |  |
|  | Treatment | Male | 10 | 6 | 6 |  |  |
| T1 (F1T1) | Control | Female | 10 | 6 | 6 |  |  |
|  | Control | Male | 10 | 6 | 5 | ML145 | The mt-rRNA content of the sample exceeded 10% of the total amount of reads |
|  | Treatment | Female | 10 | 6 | 6 |  |  |
|  | Treatment | Male | 10 | 6 | 6 |  |  |
| T1 (F2T1) | Control | Female | 10 | 6 | 6 |  |  |
|  | Control | Male | 10 | 6 | 6 |  |  |
|  | Treatment | Female | 10 | 6 | 6 |  |  |
|  | Treatment | Male | 10 | 6 | 6 |  |  |
