## Supplementary material for "Persistent maternal memory, transient offspring effects: the temporal and sex-specific architecture of heritable heat-stress responses in zebrafish": SupplementaryFile08.TableS8_Outliers_Oocytes.docx

**Supplementary File 08.**

**Table S8. An overview of the sequenced unfertilized oocyte samples.**

Before the differential gene expression analysis, the RNA-seq data obtained were investigated for the presence of any sequencing quality issues. If any of those were identified, the samples were not included in the further analysis. Samples that were excluded based on the quality control results before the sequencing (low RIN value or signs of rRNA degradation) were not included in the lists below (except for the cases where all samples from the same mother were excluded). The table provides a detailed overview of the samples that are included and excluded (as well as the reasoning behind this).

| Timepoint (Population) | Group | Mother ID | Samples N | | | Excluded samples | Reason for exclusion |
| --- | --- | --- | --- | --- | --- | --- | --- |
|  |  |  | extracted | sequenced | analyzed |  |  |
| T1 (F1T1) | Treatment | D1 | 11 | 9 | 0 | All d1 | High number of multimapper genes in many samples; the amount of reads in many samples is a few times higher than in all other samples from other mothers (can be up to 20x times higher) |
|  | Treatment | D2 | 14 | 12 | 12 |  |  |
|  | Treatment | D3 | 17 | 16 | 11 | d3_18  d3_5  d3_12  d3_14  d3_15 | Separate clustering from other samples derived from the same mother on the sample-to-sample distance matrix. |
|  | Control | D4 | 19 | 12 | 9 | d4_2  d4_17  d4_10 | d4_2, d4_17: the mt-rRNA content of the sample exceeded 10% of the total amount of reads  d4_10: Low number of mapped reads (< 10 Mio) |
|  | Control | D5 | 16 | 15 | 0 | All d5 | Very distant clustering from all other samples within the same timepoint; unexplained upregulation of all genes in comparison to the other control mothers (suspected, but not confirmed issues with UMIs) |
|  | Control | D6 | 14 | 0 | 0 | All d6 | All samples had low RIN values and signs of rRNA degradation (not sequenced) |
|  | Treatment | D7 | 20 | 18 | 18 |  |  |
|  | Treatment | D8 | 19 | 18 | 18 |  |  |
|  | Treatment | D9 | 19 | 17 | 17 |  |  |
|  | Control | D10 | 19 | 15 | 14 | d10_19 | Distant from other samples derived from the same mother on the PCA plot; not transparent during the elution step |
|  | Control | D11 | 19 | 18 | 17 | d11_1 | Low number of mapped reads (< 20 Mio) |
|  | Control | D12 | 19 | 8 | 8 |  |  |
| T2 (F1T2) | Treatment | D13 | 18 | 12 | 12 |  |  |
|  | Treatment | D14 | 19 | 12 | 12 |  |  |
|  | Treatment | D15 | 17 | 0 | 0 | All d15 | All samples had low RIN values and signs of rRNA degradation (not sequenced) |
|  | Control | D16 | 20 | 12 | 12 |  |  |
|  | Control | D17 | 20 | 12 | 9 | d17_11  d17_9  d17_15 | The mt-rRNA content of the sample exceeded 10% of the total amount of reads |
|  | Control | D18 | 18 | 12 | 10 | d18_10  d18_2 | The mt-rRNA content of the sample exceeded 10% of the total amount of reads |
|  | Treatment | D19 | 19 | 12 | 11 | d19_11 | The mt-rRNA content of the sample exceeded 10% of the total amount of reads |
|  | Treatment | D20 | 19 | 12 | 12 |  |  |
|  | Treatment | D21 | 19 | 12 | 11 | d21_6 | The mt-rRNA content of the sample exceeded 10% of the total amount of reads |
|  | Control | D22 | 19 | 12 | 12 |  |  |
|  | Control | D23 | 19 | 12 | 12 |  |  |
|  | Control | D24 | 17 | 12 | 12 |  |  |
| T3 (F1T3) | Treatment | D25 | 19 | 12 | 12 |  |  |
|  | Treatment | D26 | 19 | 12 | 11 | d26_19 | The mt-rRNA content of the sample exceeded 10% of the total amount of reads |
|  | Treatment | D27 | 20 | 12 | 12 |  |  |
|  | Control | D28 | 17 | 12 | 0 | All d28 | Poor quality oocytes during sampling (a lot of damaged oocytes released by the mother); mt-rRNA content exceeding 10% of the total amount of reads for many samples |
|  | Control | D29 | 16 | 12 | 12 |  |  |
|  | Control | D30 | 19 | 12 | 12 |  |  |
|  | Treatment | D31 | 19 | 12 | 12 |  |  |
|  | Treatment | D32 | 19 | 12 | 12 |  |  |
|  | Treatment | D33 | 19 | 12 | 12 |  |  |
|  | Control | D34 | 19 | 12 | 12 |  |  |
|  | Control | D35 | 17 | 12 | 12 |  |  |
|  | Control | D36 | 20 | 12 | 12 |  |  |
| T1 (F2T1) | Treatment | D58 | 20 | 12 | 12 |  |  |
|  | Treatment | D59 | 19 | 12 | 12 |  |  |
|  | Treatment | D60 | 20 | 12 | 12 |  |  |
|  | Control | D61 | 20 | 12 | 12 |  |  |
|  | Control | D62 | 19 | 12 | 12 |  |  |
|  | Control | D63 | 20 | 12 | 12 |  |  |
|  | Treatment | D64 | 20 | 12 | 12 |  |  |
|  | Treatment | D65 | 20 | 12 | 12 |  |  |
|  | Treatment | D66 | 20 | 12 | 12 |  |  |
|  | Control | D67 | 19 | 12 | 12 |  |  |
|  | Control | D68 | 19 | 12 | 12 |  |  |
|  | Control | D69 | 19 | 12 | 12 |  |  |
