## Supplementary material for "Persistent maternal memory, transient offspring effects: the temporal and sex-specific architecture of heritable heat-stress responses in zebrafish": SupplementaryFile14.TableS14_GeneOverlap_matrix.docx

**Supplementary File 14.**

**Table S14. Gene overlap matrix of significantly differentially expressed genes in oocytes and livers across time points and generations.**

The matrix shows the number of significantly differentially expressed genes shared between each pair of gene sets across oocytes and liver tissue (females and males), time points, and generations. Gene sets were generated by extracting significantly up- and downregulated genes from DESeq2 results using tissue-specific thresholds. For liver samples, genes with a log₂ fold change ≥ 1.5 and an adjusted *P*-value ≤ 0.05 were included. For oocyte samples, genes with a log₂ fold change ≥ 2 and an adjusted *P*-value ≤ 0.05 were included.

|  | | | F0 | | | | | | F1 | | | | | F2 | | |
| --- | --- | --- | --- | --- | --- | --- | --- | --- | --- | --- | --- | --- | --- | --- | --- | --- |
|  |  |  | T1 | | T2 | | T3 | | T1 | | | T2 | T3 | T1 | | |
|  |  |  | Male | Female | Male | Female | Male | Female | Oocyte | Male | Female | Oocyte | Oocyte | Oocyte | Male | Female |
| F0 | T1 | Male | **127** | 0 | 10 | 1 | 1 | 11 | 5 | 5 | 0 | 1 | 0 | 0 | 7 | 4 |
|  |  | Female | 0 | **5** | 1 | 0 | 2 | 1 | 0 | 0 | 0 | 0 | 0 | 0 | 0 | 1 |
|  | T2 | Male | 10 | 1 | **133** | 0 | 4 | 9 | 3 | 7 | 5 | 3 | 0 | 1 | 4 | 4 |
|  |  | Female | 1 | 0 | 0 | **5** | 0 | 1 | 0 | 0 | 0 | 0 | 0 | 0 | 0 | 0 |
|  | T3 | Male | 1 | 2 | 4 | 0 | **28** | 4 | 3 | 8 | 0 | 1 | 2 | 1 | 11 | 1 |
|  |  | Female | 11 | 1 | 9 | 1 | 4 | **183** | 4 | 14 | 4 | 4 | 2 | 2 | 9 | 2 |
| F1 | T1 | Oocyte | 5 | 0 | 3 | 0 | 3 | 4 | **427** | 1 | 1 | 57 | 49 | 27 | 4 | 0 |
|  |  | Male | 5 | 0 | 7 | 0 | 8 | 14 | 1 | **97** | 1 | 2 | 0 | 2 | 29 | 2 |
|  |  | Female | 0 | 0 | 5 | 0 | 0 | 4 | 1 | 1 | **33** | 1 | 0 | 1 | 1 | 0 |
|  | T2 | Oocyte | 1 | 0 | 3 | 0 | 1 | 4 | 57 | 2 | 1 | **187** | 40 | 25 | 2 | 0 |
|  | T3 | Oocyte | 0 | 0 | 0 | 0 | 2 | 2 | 49 | 0 | 0 | 40 | **215** | 37 | 2 | 1 |
| F2 | T1 | Oocyte | 0 | 0 | 1 | 0 | 1 | 2 | 27 | 2 | 1 | 25 | 37 | **163** | 1 | 1 |
|  |  | Male | 7 | 0 | 4 | 0 | 11 | 9 | 4 | 29 | 1 | 2 | 2 | 1 | **90** | 0 |
|  |  | Female | 4 | 1 | 4 | 0 | 1 | 2 | 0 | 2 | 0 | 0 | 1 | 0 | 1 | **44** |

**Note**: F0, F1, F2 – generations used in the study; T1, T2, T3 – experimental timepoints used in the study
