## Supplementary figures and images for "Persistent maternal memory, transient offspring effects: the temporal and sex-specific architecture of heritable heat-stress responses in zebrafish"

### SupplementaryFile15.FigureS1_Correlation_plots-Livers-Oocytes.png

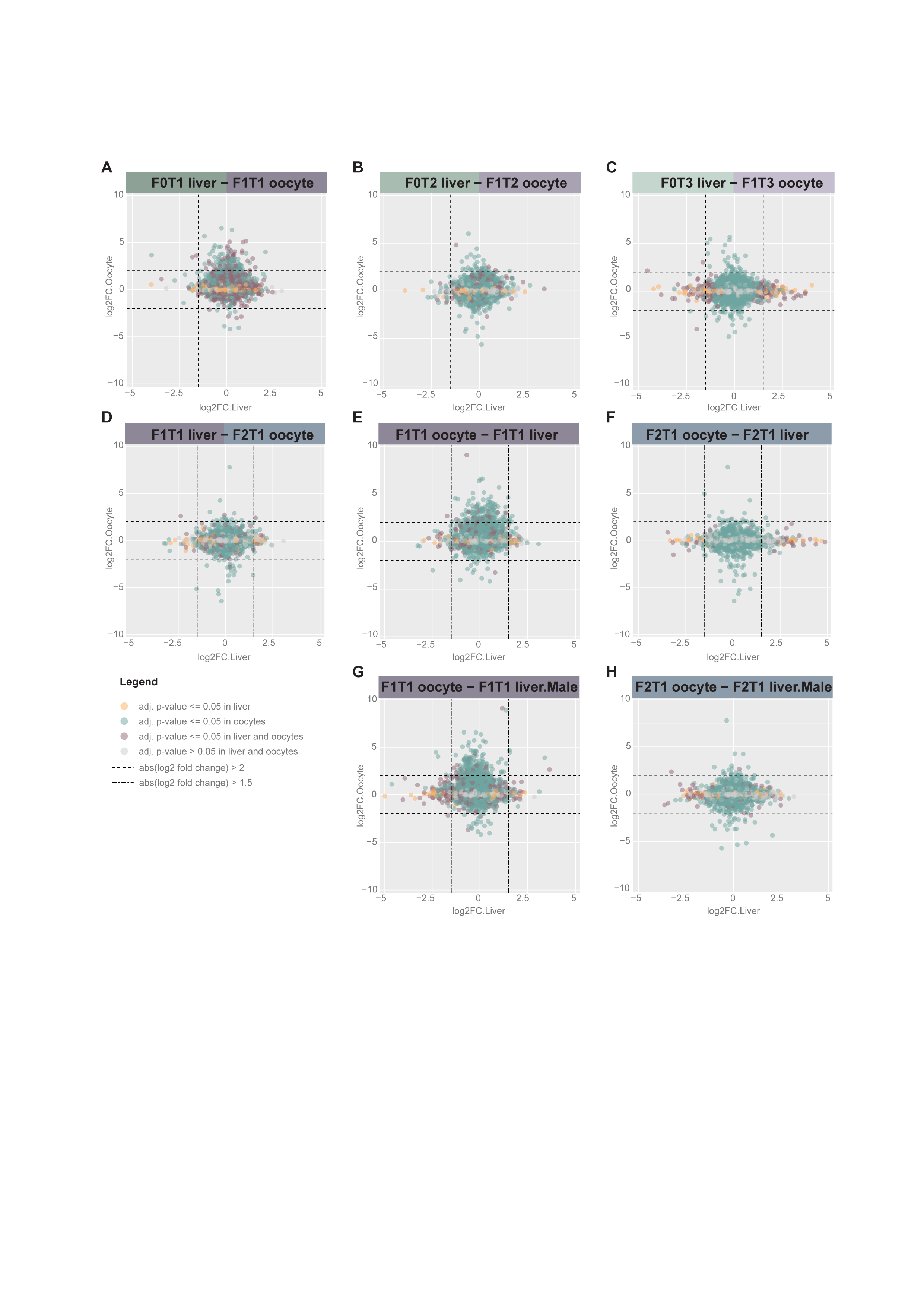
